# Speciation as a Sieve for Ancestral Polymorphism

**DOI:** 10.1101/155176

**Authors:** Rafael F. Guerrero, Matthew W. Hahn

## Abstract

Studying the process of speciation using patterns of genomic divergence between species requires that we understand the determinants of genetic diversity within species. Because sequence diversity in an ancestral population determines the starting point from which divergent populations accumulate differences (Gillespie & Langley 1979), any evolutionary forces that shape diversity within species can have a large impact on measures of divergence between species. These forces include those both decreasing (e.g., selective sweeps (Begun *et al.* 2007; Cruickshank & Hahn 2014) or background selection (Phung *et al.* 2016)) and increasing variation (e.g., balancing selection (Charlesworth 2006)). Selection can increase diversity by favoring the maintenance of polymorphism via overdominance, frequency dependence, and heterogeneous selection. Nevertheless, balanced polymorphisms are considered rare in nature, and such loci are often overlooked as major contributors to genome-wide variation in levels of sequence diversity and divergence.

Here, we argue that speciation can act as a “sieve” that will reveal otherwise elusive balancing selection by sorting ancestral balanced polymorphisms unequally across descendent lineages. By sorting alternative alleles between different species, this process uncovers the existence of balancing selection in the ancestral population as regions of higher-than-expected divergence. Sorted ancestral polymorphism may also be responsible for many of the observed peaks of genomic divergence between closely related taxa, confounding studies of differential gene flow and “islands of speciation.”

## The effect of balancing selection on genetic diversity

Balancing selection encompasses various, rather disparate, mechanisms that favor the maintenance of polymorphism. Most forms of balancing selection involve heterogeneous or variable selective forces: polymorphism is favored when selection varies across space or time (reviewed in Felsenstein (1976)), between the sexes (reviewed in Otto *et al.* (2011)), or as a function of allele frequency (negative frequency-dependent selection; reviewed in Ayala & Campbell (1974)). Balanced polymorphisms can also occur under constant selective pressures when heterozygotes have a fitness advantage over homozygotes (overdominance; (Wright 1931)).

Despite clear differences in their underlying mechanisms, all forms of balancing selection have qualitatively similar long-term effects on linked neutral variation. Genomic regions closely linked to loci under balancing selection are expected to show increased divergence between the allelic classes defined by the balanced polymorphism. This increase in divergence—the extent of which depends on population size, recombination rate, and strength of balancing selection— results from a population subdivision imposed by the balanced polymorphism. When two (or more) balanced alleles persist for a long time in a population, closely linked regions tend to accumulate differences between the allelic classes. Thus, the genealogies of samples linked to balanced polymorphisms resemble those of structured populations, with allelic classes acting as subpopulations and recombination allowing “gene flow” between these subpopulations. However, when the balanced alleles at a locus are not known, samples cannot be partitioned by allelic class. This makes it difficult to quantify the increased divergence between allelic classes directly (for example, using the statistic *F*_ST_). Instead, the only observable signals of balanced polymorphism are an excess of intermediate frequency alleles (detected using Tajima’s *D*), or overall increases in diversity (detected using the average number of pairwise differences, or π). Unfortunately, the large amount of variance in both *D* and π associated with even neutrally evolving loci means that loci under balancing selection are hard to detect (Simonsen *et al.* 1995).

The potential difficulty in detecting balanced polymorphism is somewhat alleviated in cases of local adaptation. Because alternative balanced alleles differ in frequency between the populations where they are individually advantageous, measures of differentiation between populations can be used as a proxy for divergence between allelic classes. Moreover, actual population structure can lower the effective recombination rate between allelic classes, exacerbating their divergence. Perhaps due to its increased detectability, spatially varying selection is reported in natural populations much more frequently than other forms of balancing selection (Asthana *et al.* 2005; Charlesworth 2006; Fan *et al.* 2016). Indeed, while overdominance and negative frequency-dependence continue to be considered rare, local adaptation is considered pervasive, and is even a compulsory first step in some models of speciation (Nosil 2012).

## The sieve: ancestral lineage sorting after speciation

Consider a simple case of allopatric speciation: a single population is split in two by vicariance (*e.g.*, the rise of a mountain range or the construction of a thousand-mile wall). For neutral biallelic polymorphisms in the ancestral population, drift will fix alternative alleles in the two nascent species at half of all such loci (the rest of the time both species will fix the same allele). In the presence of balancing selection, expectations can differ. Balancing selection may favor the maintenance of polymorphism in both nascent species (resulting in ‘trans-specific polymorphism’; (Muirhead *et al.* 2002)), or increase the chance that alternative alleles are fixed. For instance, when selection varies across space, a geographic barrier is likely to create two unequal ranges (*i.e.*, areas with different proportions of habitats driving local adaptation)—these unequal habitat ranges may favor dramatically different equilibrium frequencies at the locally adapted loci in the nascent species. This could result in selection favoring the fixation of opposite alleles in each species, sieving ancestral alleles in the descendant lineages (Figure 1A).

**Figure 1.**
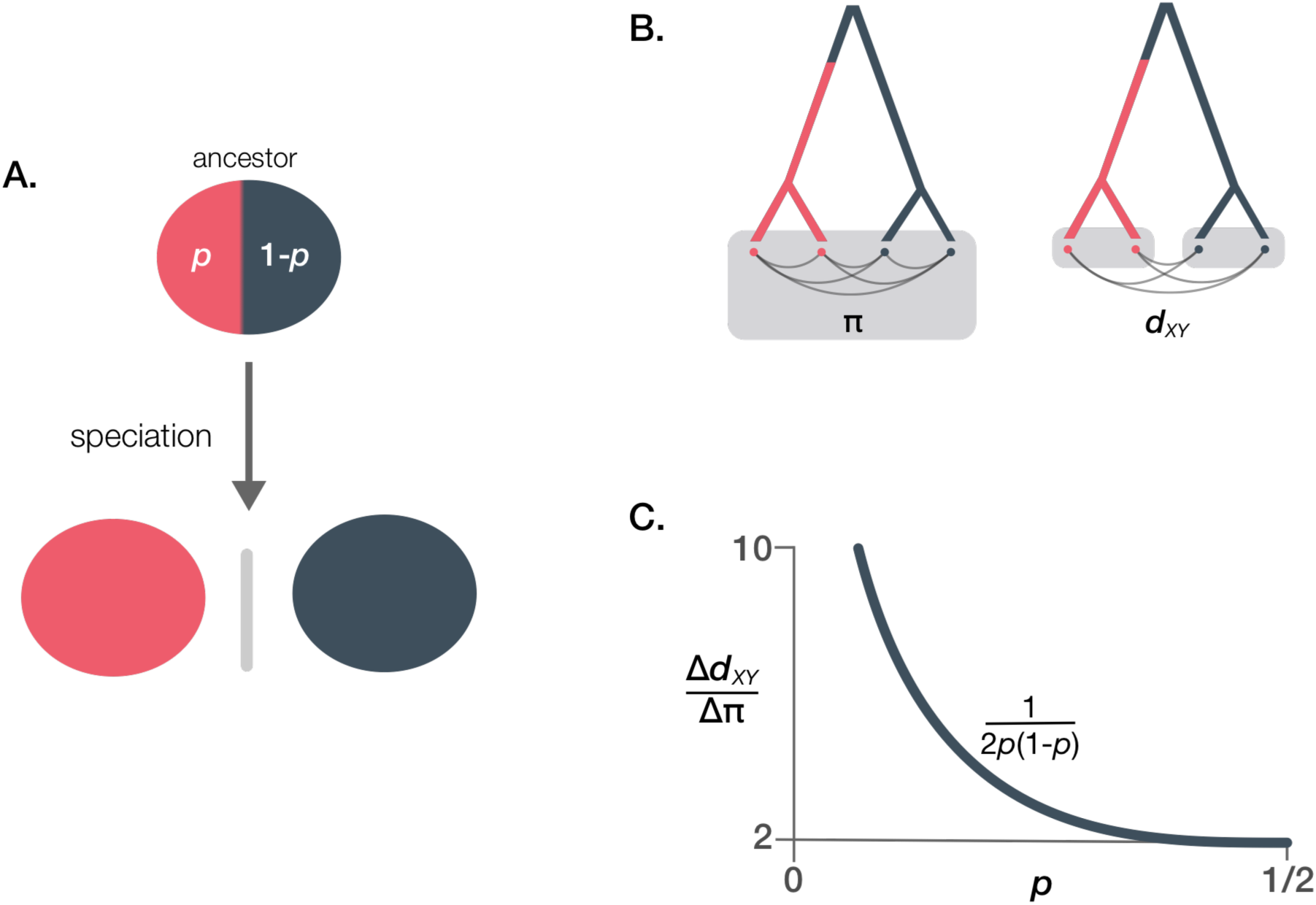
A) Schematic of a sieved polymorphism. In the ancestor, two alleles are balanced at frequencies *p* and 1-*p*, and after speciation different alleles fix in descendant lineages. B) The increased power to detect balancing selection stems from the partitioning of the ancestral population imposed by the sieve. While π is a measure of all pairwise distances in a population (left), *d_XY_* compares only samples from different allelic classes (right). C) The ratio of the effect of a balanced polymorphism on diversity (Δπ = π - π_o_, where π_o_ is the baseline diversity) and on divergence (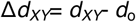, and *d*_o_ = π_o_) increases as the minor allele becomes rare. As the time since the origin of the polymorphism increases, the ratio converges to the inverse of the expected heterozygosity, *i.e*.,1/2*p*(1-*p*).

Sieved balanced polymorphisms carry the signature of selection in the form of increased sequence divergence between the descendant species. Immediately after the split, the level of genetic divergence is largely determined by the diversity present in the ancestral population. Formally, this can be seen in the expectation of absolute divergence: E(*d_XY_*)=2μ*t*+θ_Anc_ (Gillespie & Langley 1979). As the time since the species split (*t*) approaches zero, E(*d_XY_*) becomes approximately equal to the ancestral level of diversity, θ_Anc_ (=4*N*_e_μ for diploids, where *N*_e_ is the effective population size and μ is the neutral mutation rate). Initial levels of divergence are therefore strongly affected by forces that affect levels of ancestral neutral diversity, such as balancing selection. In regions linked to sieved balanced polymorphism, measures of divergence between species (such as *d_XY_* or *F*_ST_) reflect the divergence accumulated between allelic classes both in the ancestor and since the lineages split, so sieved polymorphisms maintained in the ancestor for a long time can appear as regions of elevated divergence between nascent species.

Interestingly, this implies that balancing selection could be more readily detected after speciation (as a sieved polymorphism with elevated *d_XY_*) than in the ancestor when the causal alleles are unknown (using *D* or π; Figure 1B). Indeed, the signature of balancing selection is expected to be twice as strong on *d_XY_* than on π at loci with alleles maintained at equal frequencies in the ancestor (Figure 1C; see Supplement). As either allele becomes rarer, the effect is more severe: when the minor allele frequency was 10% in the ancestor, the increase in *d_XY_* is expected to be almost ten times larger than the increase in π. Therefore, by separating the relevant haplotypes, speciation can dramatically increase our power to find loci under balancing selection.

## Relevance of sieved polymorphisms during recent genomic divergence

The prevalence of sieved polymorphisms in nature is unknown (and is probably low), but the importance of this phenomenon for observed patterns of genomic divergence does not stem from its frequency. The amount of balanced polymorphism sieved by speciation is proportional to the fraction of loci under balancing selection in the ancestor and the probability of fixing alternative alleles at those loci (which depends on the mode of selection operating at each locus). If balancing selection is as rare as usually assumed (Charlesworth 2006), sieved polymorphisms are likely to be uncommon and would not drastically elevate average levels of divergence. Instead, the few instances of sieved polymorphisms will have subtle but significant repercussions: these regions will tend to appear at the top of the distribution of divergence across the genome, ‘fattening’ its upper tail and potentially affecting further inferences. The potential consequences for the distribution of divergence depend not only on the fraction of loci sieved, but also on parameters specific to each polymorphism (namely, its age, strength of selection, and recombination rate). If, for instance, a genome carries only one sieved polymorphism, it will likely appear as a divergence outlier. On the other hand, a higher fraction of sieved regions—caused, for instance, by numerous locally adapted alleles differentiated between populations prior to speciation—will considerably fatten the upper tail in the divergence distribution. Such an observation could be interpreted as evidence of a period of differential gene flow following an initial split (*cf*. Yang et al. (2017)). This latter case highlights the fact that sieved polymorphism can mimic the signature of other evolutionary processes, and distinguishing among these may be challenging without additional pieces of evidence (see below).

Recent findings suggest that sieved polymorphisms do play an important role in shaping patterns of genomic divergence. Multiple studies have found regions with levels of divergence so high that differentiation at these loci almost certainly started before speciation, consistent with ancestral balanced polymorphism. In these cases, the timing of speciation—or at least a bound on the timing—can be independently estimated, highlighting the mismatch between species divergence and genetic divergence.

In the radiation of Darwin’s finches, speciation events happened roughly between 50 to 500 thousand years ago (Lamichhaney *et al.* 2015), yet some genomic regions seem to have started differentiating well before then (up to one million years ago; (Han *et al.* 2017)). At least two of these genomic regions, linked to loci associated with beak shape and size (genes *ALX1* and *HMGA2*), are likely sieved polymorphisms. Across the nine species of tree and ground finches studied, these loci have two distinct haplotype classes (*i.e.*, there is high divergence between classes and reduced divergence within class, even between species) that are responsible for marked phenotypic differences (blunt vs. pointed beaks, and small vs. large beaks). As a result, species pairs that have fixed different haplotype classes show ‘islands of divergence’ at these loci. As hypothesized by Han *et al.* (2017), however, this beak polymorphism was probably balanced in the ancestor (perhaps under negative frequency-dependent selection) and was later sieved across the Galapagos (Han *et al.* 2017).

In the freshwater threespine sticklebacks of western North America, the *Eda* locus represents a clear example of how ancestral polymorphisms can emerge as conspicuous peaks of genomic divergence. Polymorphism at *Eda* has been maintained in marine populations for approximately two million years, and the minor allele has been selected repeatedly during colonization events of glacier lakes around ten thousand years ago (Colosimo *et al.* 2005). Expectedly, *Eda* shows dramatic differentiation between marine and freshwater populations, but most of this divergence happened before the invasion of the glacier lakes and is unrelated to recent processes.

Other types of polymorphism can also be sieved. Among these, chromosome rearrangements (*e.g.*, inversions, fusions) are of special interest for their role during local adaptation. Rearrangements can evolve by capturing locally adapted alleles (Kirkpatrick & Barton 2006; Guerrero & Kirkpatrick 2014), and established rearrangements can promote further local adaptation (Navarro & Barton 2003). Moreover, balanced rearrangements (especially inversions) are usually conspicuous in population genomic data (*e.g.*, (Cheng *et al.* 2012; Kapun *et al.* 2016)), as they typically cause a dramatic reduction in recombination, which in turn leads to much stronger population subdivision compared to other balanced polymorphisms (Guerrero *et al.* 2012; Guerrero & Kirkpatrick 2014). Due to their role in local adaptation and their large genomic footprint, rearrangements are thought to be key players in the buildup of differentiation that can lead to speciation. Some chromosome inversions have in fact been linked to speciation processes (*e.g.*, in cactophilic *Drosophila* (Lohse *et al.* 2015), *Mimulus* (Fishman *et al.* 2013)). In other cases, however, locally adapted inversions are maintained as polymorphisms within a species or as trans-specific polymorphisms – without being involved in speciation. In the *Anopheles gambiae* species complex, inversion *2La* arose in the ancestor of six species (well before the most recent speciation events) and it is still polymorphic in two of these (Fontaine *et al.* 2015). This inversion has been sieved at least two times, such that comparisons between species fixed for alternative arrangements show increased divergence across many megabases of sequence.

## Implications for inferences of speciation islands and speciation-with-gene-flow

Many of the patterns described above for sieved polymorphisms mimic predictions of models of speciation with gene flow. In the most common version of this model (which is conceptually similar to sympatric and parapatric speciation models; reviewed in (Bush 1975; Via 2001)), populations have uninterrupted exchange of migrants, but achieve total reproductive isolation gradually by the accumulation of locally adapted loci that limit effective migration (*cf.* Charlesworth *et al.* (1997)). At the genomic level, differential effective migration is expected to leave a clear signature: regions linked to loci under divergent selection (*a.k.a.* local adaptation) accumulate higher divergence compared to the rest of the genome, appearing as ‘genomic islands of speciation’ (Turner *et al.* 2005). Variation in divergence across the genome has therefore been attributed to differential migration among loci, with individual loci showing much higher levels of divergence implicated as being causal in the speciation process. Several studies initially reported finding such islands using relative measures of divergence (such as *F*_ST_), which can be affected by selection in the sampled populations (*e.g.*, (Turner *et al.* 2005; Geraldes *et al.* 2011; Ellegren *et al.* 2012)). Because absolute measures of divergence (such as *d_XY_*) are unaffected by current levels of polymorphism, it was suggested that these would be preferred in identifying regions that are truly resistant to introgression (Cruickshank & Hahn 2014). It has therefore become more common to search for islands using *d_XY_* and related statistics, and some researchers have reported finding these important loci (*e.g.*, (Malinsky *et al.* 2015; Marques *et al.* 2016)).

Using absolute measures of divergence does not obviate the problem of variation in levels of diversity in the ancestral population. As discussed above, regions of elevated *d_XY_* can be produced by balanced polymorphisms in ancestral populations, and variance in levels of divergence across the genome can be driven by variance in diversity in these ancestral populations. It is simply not true that all loci start out equally diverged at speciation, or that differential migration is the only force that can produce variation in *d_XY_* beyond that expected from neutral coalescent variation in the ancestor.

How would one distinguish between true islands and sieved balanced polymorphisms? One commonality shared by the clearest examples of sieved polymorphisms given above is that independent estimates exist for the earliest time when speciation could have started. Glacial lakes that could not have existed prior to the retreat of the glaciers, radiations onto geological features (such as oceanic islands) that recently appeared on the landscape, or simply the date of an earlier divergence from a more distantly related species, all limit the maximum time pairs of focal species could have been separated. Given such limits, we can then contrast hypotheses of speciation-with-gene-flow with those involving sieved polymorphisms (Figure 2). In fact, it takes quite a long time for loci resistant to gene flow to appear as divergence outliers under models of speciation-with-gene-flow (Figure 2A; also see Fig. B1 in Cruickshank and Hahn 2014). By contrast, sieved balanced polymorphisms can be detected almost immediately by both relative and absolute measures of divergence, and ironically these signals are stronger in *F*_ST_ than *d_XY_* (Figure 2B).

**Figure 2.**
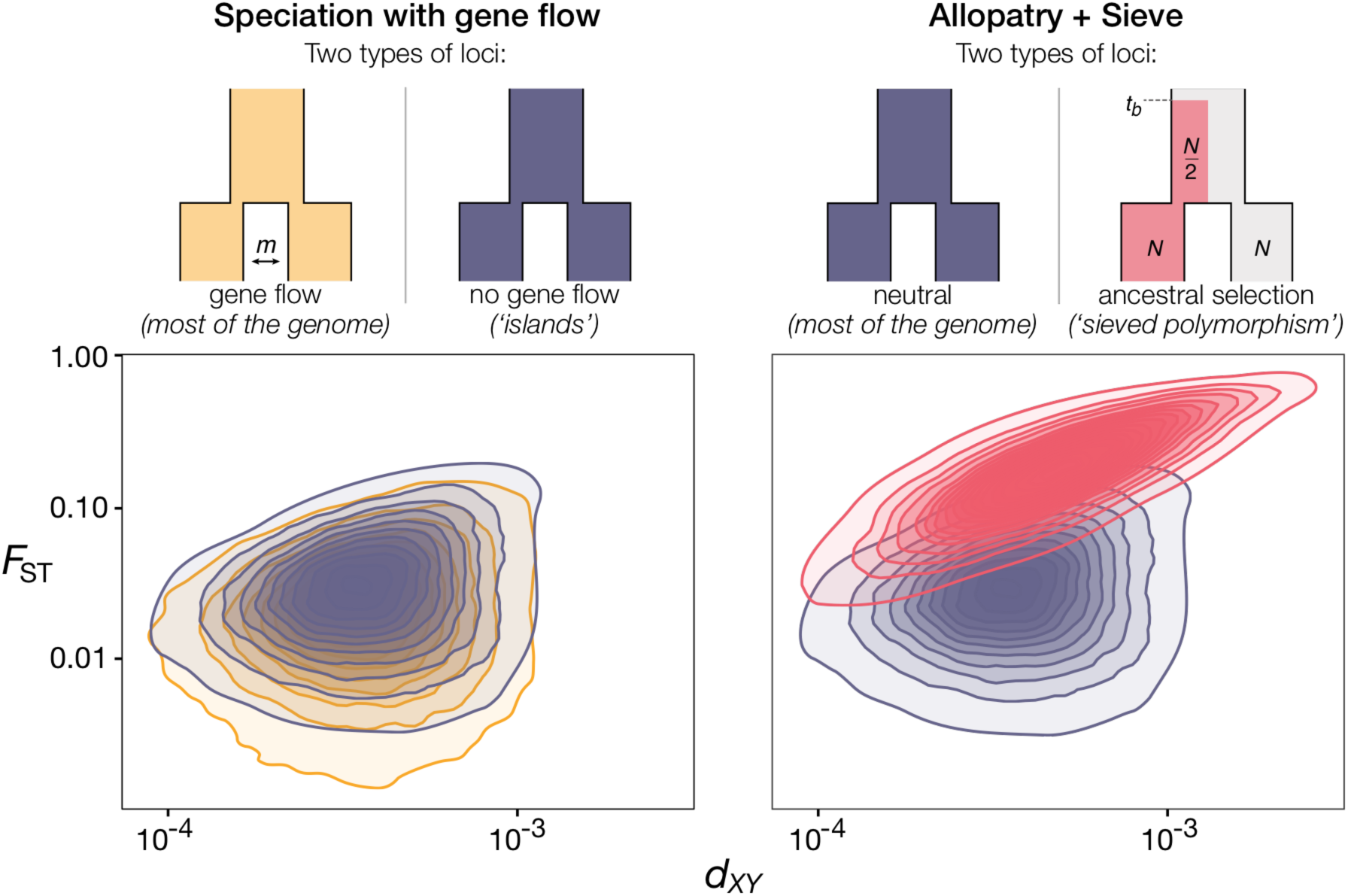
Distributions of absolute and relative divergence (*d_XY_* and *F*_ST_) for genomes under two scenarios of recent speciation (1000 generations ago, population size *N*=10^4^ for each species). In each scenario, genomes have two types of regions. On the left, regions experience differential gene flow since the split: while in most of the genome (in light orange, *m*=0.0001) gene flow prevents divergence, in “speciation islands” (in purple, *m*=0) there is a slight increase in *F*_ST_. On the right, there is no gene flow after speciation, but some regions are tightly linked to a sieved polymorphism (in pink; balanced locus is at *r*=10^-5^ from the simulated region, originated *t_b_* = 10^4^ generations ago, stable at frequency of ½ in ancestor, alternative alleles are fixed in descendants). We simulated the genealogy for a sample of 20 chromosomes drawn from each species (10^5^ coalescent simulations for each type of genomic region, a 10 Kb non-recombining segment with μ=10^-8^).

We can apply these ideas to assess the likely causes of “islands of divergence” in previously published examples. In the cichilds of Lake Massoko, for instance, levels of divergence might stem from ancestral polymorphism (Malinsky *et al.* 2015). The speciation process in the focal pair started some 3500 generations ago (Malinsky *et al.* 2015), which would allow enough time for the accumulation of divergence on the order of *d_XY_*=1.05x10^-4^ (assuming μ=1.5x10^-8^ and θ_Anc_=0). However, *d_XY_* observed in the most highly diverged regions is considerably higher than this expectation (mean *d_XY_*=9x10^-4^; https://twitter.com/millanek1/status/758209899964862465), suggesting that a large fraction of the observed differentiation is due to ancestral diversity. This rough calculation ignores many factors, such as variation in mutation rate, that can contribute to the observed patterns. However, it allows us to emphasize that the genomes of extant populations give us a glimpse into ancestral processes that transcend the most recent speciation event.

Similarly, in the threespine stickleback of Lake Constance, subspecies show several regions of divergence that most likely predate the current process of local adaptation (which started about 150 generations ago), and for which standing variation has been invoked as a probable source (Marques *et al.* 2016). In this case, high differentiation was inferred in 37 genomic regions based on allele frequency differences among populations (using SNPs obtained via RAD-seq). The observed diversity levels in these regions are not significantly reduced, suggesting that—while current selection may be driving allele frequency divergence—the accumulation of divergent SNPs is not the result of a recent sweep. Rather, many of these “islands” are likely ancestral balanced haplotypes currently being sorted.

Confounding sieved polymorphism with islands of speciation can lead to an additional erroneous inference. The observation of large amounts of variation in divergence time among loci may lead to the conclusion that gene flow has occurred, when none has. To some extent, these inferences follow from the observation of islands—if there are loci resistant to gene flow, then it follows that there must have been gene flow. But this false signal can also affect methods for inferring gene flow that assume that there is no selection, and therefore interpret the excess variance observed as due to migration. It has recently been recognized that modeling the effects of selection on the levels of sampled polymorphism is important in controlling such false positives (Roux *et al.* 2016). The implication here is that variation in levels of ancestral polymorphism must also be considered, since they can lead to false inference of gene flow and current selection (*i.e.*, labeling sieved regions, which may be neutral now, as resistant to gene flow).

## Conclusions

Sieved polymorphism—in conjunction with factors such as population structure, assortative mating, background selection, or variation in mutation and recombination rates—contributes to heterogeneity in genomic divergence levels. Due to the complexity of the divergence distribution, inferences that rely solely on its outliers (*e.g.*, taking an arbitrary upper quantile of *d_XY_* as speciation islands) can yield misleading results by selecting regions, such as sieved polymorphisms, that are unrelated to speciation. Model-based analyses are necessary but not sufficient, because extremely similar patterns of genomic divergence can be generated by alternative models of speciation. In fact, biologically significant differences between speciation models are occasionally irrelevant from a theoretical standpoint (*e.g.*, cessation of gene flow is modeled identically [*m*=0] regardless of the mechanism behind it, whether due to hybrid inviability, a geographic barrier, or other). For this reason, independent lines of evidence are critical to disentangle the multiple forces at play. For instance, having a lower bound on the time since speciation can allow us to determine how much ancestral polymorphism is expected, since its effect is strongest in recent speciation events (*i.e.*, *t* < 2*N*_e_, when θ_Anc_ accounts for more than half of E(*d_XY_*)). Using current levels of polymorphism as a proxy for ancestral levels in model-based analyses will also be a useful starting point in trying to understand the causes of variation in divergence levels. Finally, a search for the signals of a speciation sieve applied across the many new whole-genome datasets being produced may cause us to reconsider the frequency with which balancing selection occurs.

